# Landscape scale terrestrial factors are also vital in shaping Odonata diversity of watercourses

**DOI:** 10.1101/724476

**Authors:** H. Beáta Nagy, Zoltán László, Flóra Szabó, Lilla Szőcs, György Dévai, Béla Tóthmérész

## Abstract

Habitat loss and fragmentation causes decline of insect populations. Odonata (both dragonflies and damselflies) are especially threatened, because they are notably influenced by both aquatic and terrestrial environment. We explored the relative importance of local and landscape variables for Odonata assemblages (species richness, assemblage composition, population abundance) revealing differences in the sensitivity of Zygoptera and Anisoptera on the selected variables. Our study took two years and was placed along 11 lowland watercourses. We sampled the specimens using 500 m long transects from May to September. Landscape variables (length of watercourses, forest patch proportion, and farmland patch size) were calculated at three scales to better account for fragmentation. Our findings show that local variables influence damselflies, but dragonflies are more sensitive to landscape variables. Damselfly’s diversity decreased with the increasing macrovegetation cover, while dragonfly’s diversity decreased with the increasing degree of land use intensification, but increased with the length of watercourses. Our findings, both on local and landscape scales demonstrated the importance of terrestrial environment on Odonata. Based on our findings we stress the importance of partial watercourse clearing, and maintenance of traditional farm management based on small parcel farming near watercourses to maintain diverse and healthy Odonata assemblages.

## INTRODUCTION

Odonata are real flagship taxa of freshwater ecosystems, and often used as indicator species to assess the quality of their close environment ^1^. Their high diversity, complex life history, the relatively rapid development and their essential role in food webs ^2,3^ make them ideal model insects for ecological surveys. Healthy aquatic habitats are crucial for the development of Odonata, but beside this, adults need resource-rich terrestrial habitats as well for maturation, feeding, resting, and mating ^4^. Furthermore, Odonata are also sensitive to the landscape composition and configuration; their sensitivity to landscape can even exceed those of water hydrography and chemistry or other local ecological parameters describing the aquatic environment ^5^. The presence and abundance of Odonata along watercourses are also affected by several conditions like water quality ^6^, competition between larvae ^7^, competition between adults ^8^, dispersal ability of adults ^9^, and the surrounding landscape ^10^.

Human-caused habitat loss and fragmentation became the major threatening factor during the last few decades for several taxa ^11^. While several studies explore the influence of habitat loss on terrestrial populations and communities ^12–14^relatively few studies focus on the relationship between landscape change and aquatic invertebrates such as Odonata. These insects are strongly connected with water bodies and regarded to be influenced mostly by the aquatic environment. However, a remarkable rise in the number of studies regarding terrestrial effects on Odonata communities have emerged during the last ten years ^10,15–17^.

The literature on Odonata-environment relationship is largely restricted to single or few species, and usually consider only a few landscape variables. The majority of the existing studies are focusing on the influence of bankside and riparian vegetation, analysing mainly the presence of buffer strips or the extent of shading canopy ^1,18–20^. Other studies address the relationship of Odonata and forests ^16,21,22^, underlying the importance of trees and shrubs for these insects. Another group of studies reveal how essential is the connectivity between water bodies ^15,23–25^. Finally, a small number of studies targets Odonata assemblages using both local and landscape variables as predictors to understand occurrence, abundance and community structure of Odonata ^4,10,26^.

The goal of our study is to explore the effect of local (i.e. aquatic) and landscape variables on Odonata assemblages along lowland watercourses in two Central-Eastern European countries. Simultaneously considering the features of local habitat and the surrounding landscape we identified the variables that were essential for the maintenance of rich Odonata assemblages. We also assessed the importance of local and landscape variables by considering the two major Odonata groups separately (Zygoptera and Anisoptera) to explore taxa-specific sensitivities to the variables considered.

The specific goals of this study were to explore the relative importance of local and landscape variables for the Odonata assemblages (species richness, assemblage composition, population abundance), and explore the potential differences in the sensitivity of two major Odonata groups (Zygoptera and Anisoptera) on the selected local and landscape variables. Our study questions were: (i) Which local biotic variables affect Odonata species diversity? (ii) Which landscape variables affect Odonata species diversity? (iii) Is there any difference in these effects regarding the two suborders?

## RESULTS

During the two years, we counted 10884 specimens belonging to 34 species (Supplementary Material, Table S1). The Zygoptera and Anisoptera abundance showed no significant difference between years (χ^2^=3.54, df=1, p=0.06, Table 1). Whereas, the Zygoptera (χ^2^=1336.2, df=10, p<0.001) and Anisoptera (χ^2^=1077.5, df=10, p<0.001) site-specific mean abundances showed significant differences.

**Table 1.** Number of observed species and specimens.

| Year/group | Zygoptera | Anisoptera | Zygoptera | Anisoptera |
| --- | --- | --- | --- | --- |
|  | species |  | specimens |  |
| 2015 | 14 | 18 | 2590 | 2642 |
| 2016 | 13 | 17 | 2901 | 2751 |
| Total | 15 | 19 | 5491 | 5393 |
| Total Odonata | 34 |  | 10884 |  |

### Local biotic variables

The water depth varied between 0.2 and 1.0 meter, with an average of 0.6 meter (± 0.2). The width of watercourses varied between 1.9 and 10.4 meter, with an average of 4.2 meter (± 2.2). We found a relatively high macrovegetation cover: it varied from 6% to 95%, with an average of 72% (± 27%). Eight sites out of 11 had higher than 75% plant cover, and only one had a lower than 10%. The percentage of banksides tree cover varied between 1.6 and 65% with an average of 37% (± 23%). The average cover of herbaceous plants was relatively high: 70% (± 18), and it varied between 41% and 98%. The average plant high of the banksides was 60 cm (± 23), varying between 24-93 cm.

We found significant negative correlation between the percentage of macrovegetation cover and Odonata diversity, with increasing surface cover the diversity of Odonata decreased (Table 2). We found this correlation to be significant for Zygoptera, but not for Anisoptera diversity (Table 2). The other five local variables (water depth, water diameter, bankside cover, bankside tree cover and height of bankside vegetation) showed no significant correlation with species diversity (Table 2).

**Table 2.** Pearson correlation coefficients between local variables and species diversity, with corresponding statistics (df = 9).

|  | Odonata |  | Zygoptera |  | Anisoptera |  |
| --- | --- | --- | --- | --- | --- | --- |
|  | r | p | r | p | r | p |
| Water diameter (m) | -0.73 | 0.01 | -0.66 | 0.03 | -0.4 | 0.22 |
| Water depth (cm) | -0.04 | 0.92 | -0.46 | 0.16 | 0.02 | 0.96 |
| Water surface cover (%) | 0.06 | 0.85 | -0.22 | 0.51 | 0.08 | 0.81 |
| Bankside cover (%) | -0.39 | 0.23 | -0.44 | 0.18 | -0.05 | 0.88 |
| Bankside tree cover (%) | 0.1 | 0.77 | -0.11 | 0.76 | -0.09 | 0.78 |
| Plant height (cm) | -0.23 | 0.49 | -0.4 | 0.22 | -0.14 | 0.68 |

### Landscape variables

The landscape diversity increased from small scale (0.91±0.21) towards intermediate (0.97±0.23) and large scales (1.11±0.19). The total length of watercourses (km) within the landscape also showed an increasing trend (small scale: 5.06±2.57 km; intermediate scale: 13.83±6.85, large scale: 42.97±14.98). The forest patch proportion increased from large (9.85±5.67) to middle (12.96±13.13) and small scale (15.87±13.91). The farmland patch size (ha) decreased only from large (25.06±11.31) to middle (15.87±7.25) and small scale (14.89±7.95). The mean distance to the nearest forest patch (m) was 174.46 (±253.45) meters.

Two variables showed significant correlation with Odonata diversity from the five tested variables on landscape scale. On one hand, the total length of watercourses at the largest scale showed significant positive correlation with the diversity of Anisoptera (Table 3). The correlation was not significant for Zygoptera (Table 3), nor for the whole Odonata populations (Table 3).

**Table 3.** Pearson correlation coefficients between landscape variables and species diversity, with corresponding statistics (df=9).

|  | Scale<br>(km) | Odonata |  | Zygoptera |  | Anisoptera |  |
| --- | --- | --- | --- | --- | --- | --- | --- |
|  |  | r | p | r | p | r | p |
| Landscape diversity | 5 | -0.01 | 0.98 | -0.17 | 0.62 | 0.14 | 0.68 |
|  | 2.5 | -0.28 | 0.41 | -0.45 | 0.17 | 0.10 | 0.78 |
|  | 1.25 | -0.15 | 0.67 | -0.47 | 0.15 | 0.01 | 0.97 |
| Length of watercourses | 5 | 0.41 | 0.21 | -0.06 | 0.86 | 0.63 | 0.04 |
|  | 2.5 | 0.23 | 0.50 | -0.02 | 0.95 | 0.42 | 0.20 |
|  | 1.25 | 0.19 | 0.57 | 0.05 | 0.88 | 0.27 | 0.43 |
| Forest patch proportion | 5 | -0.22 | 0.51 | -0.46 | 0.16 | -0.01 | 0.99 |
|  | 2.5 | -0.40 | 0.23 | -0.49 | 0.13 | -0.16 | 0.64 |
|  | 1.25 | -0.55 | 0.08 | -0.55 | 0.08 | -0.24 | 0.47 |
| Farmland patch size | 5 | -0.74 | 0.01 | -0.45 | 0.16 | -0.55 | 0.08 |
|  | 2.5 | -0.45 | 0.16 | -0.14 | 0.68 | -0.60 | 0.05 |
|  | 1.25 | -0.44 | 0.17 | 0.00 | 1.00 | -0.72 | 0.01 |
| Distance to the nearest forest patch | 1.25 | 0.39 | 0.24 | 0.07 | 0.83 | 0.23 | 0.49 |

On other hand, the farmland patch size showed significant negative correlations with species diversity at various landscape scales (Figure 2). At the largest landscape scale the correlation was significant for the whole Odonata diversity (Table 3), although it was not significant for Zygoptera (Table 3), and only marginally significant for Anisoptera (Table 3). At the smaller landscape scale (radius = 2.5 km) the correlation was only marginally significant but only for Anisoptera (Table 3). We found the same pattern also at the smallest landscape scale (radius = 1.25 km); the correlation was significant for Anisoptera (Table 3).

**Figure 2.**
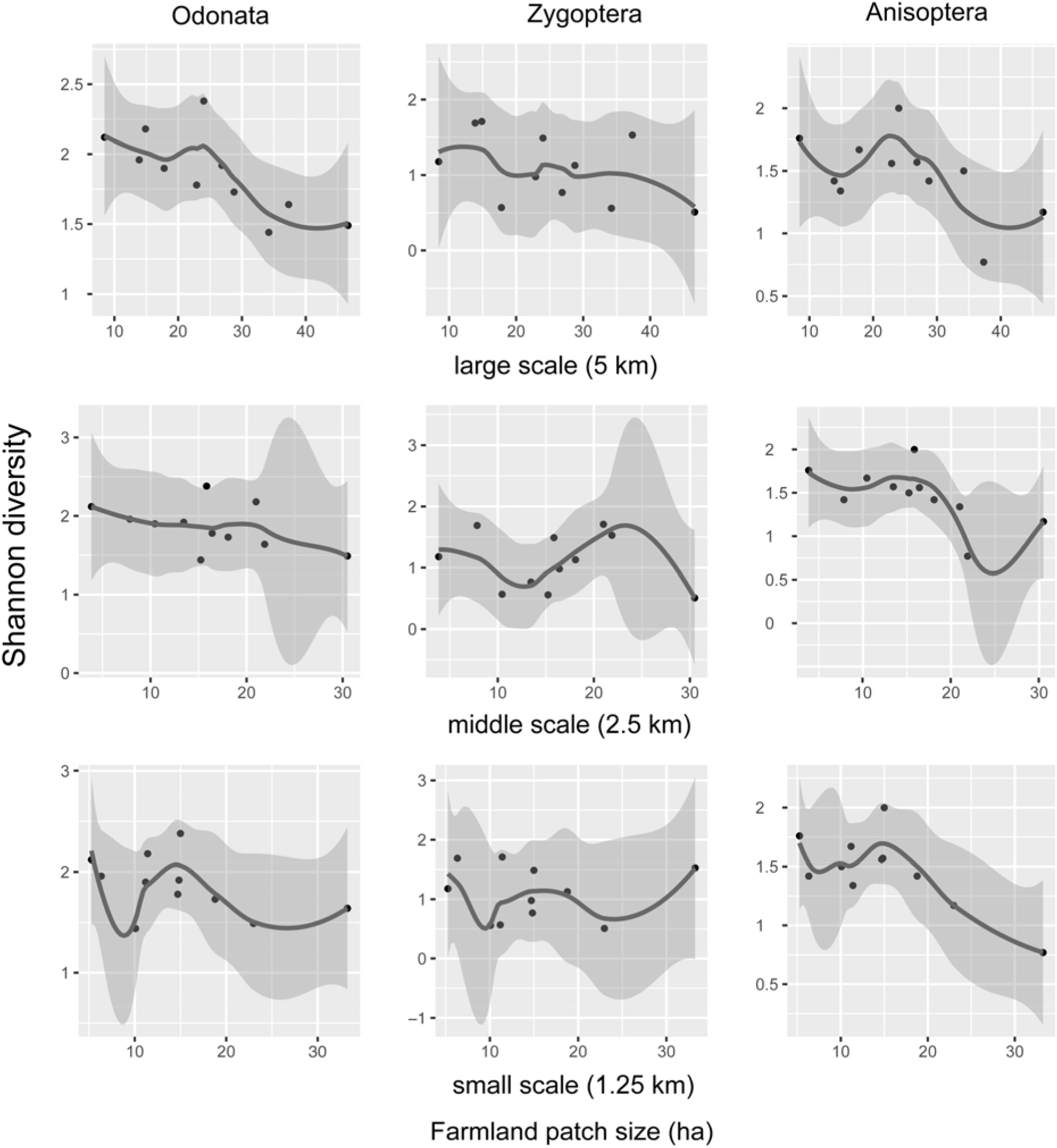
Locally weighted scatterplot smoothing curves (with 95% confidence interval around smooth – dark grey) for the relationship between farmland patch size and Odonata diversity at the studied landscape scales, considering the two suborders (Zygoptera and Anisoptera) and the Odonata assemblages.

At the smallest landscape scale (radius = 1.25 km) we also found a marginally significant correlation between the forest patch proportion and the diversity of Zygoptera (Table 3) and the diversity of whole Odonata (Table 3). Regarding the other landscape variables, we did not find any significant correlations (Table 3).

### Variable importance

The cover of emergent vegetation was the variable with the highest relative importance (both aquatic and landscape considered) in explaining the Odonata species diversity; it was followed by the farmland patch size on the 5 km scale (Table 4). The total length of watercourses on the 5 km scale had low relative importance, while the forest patch proportion on the 1.25 km scale had the lowest importance (Table 4).

**Table 4.**
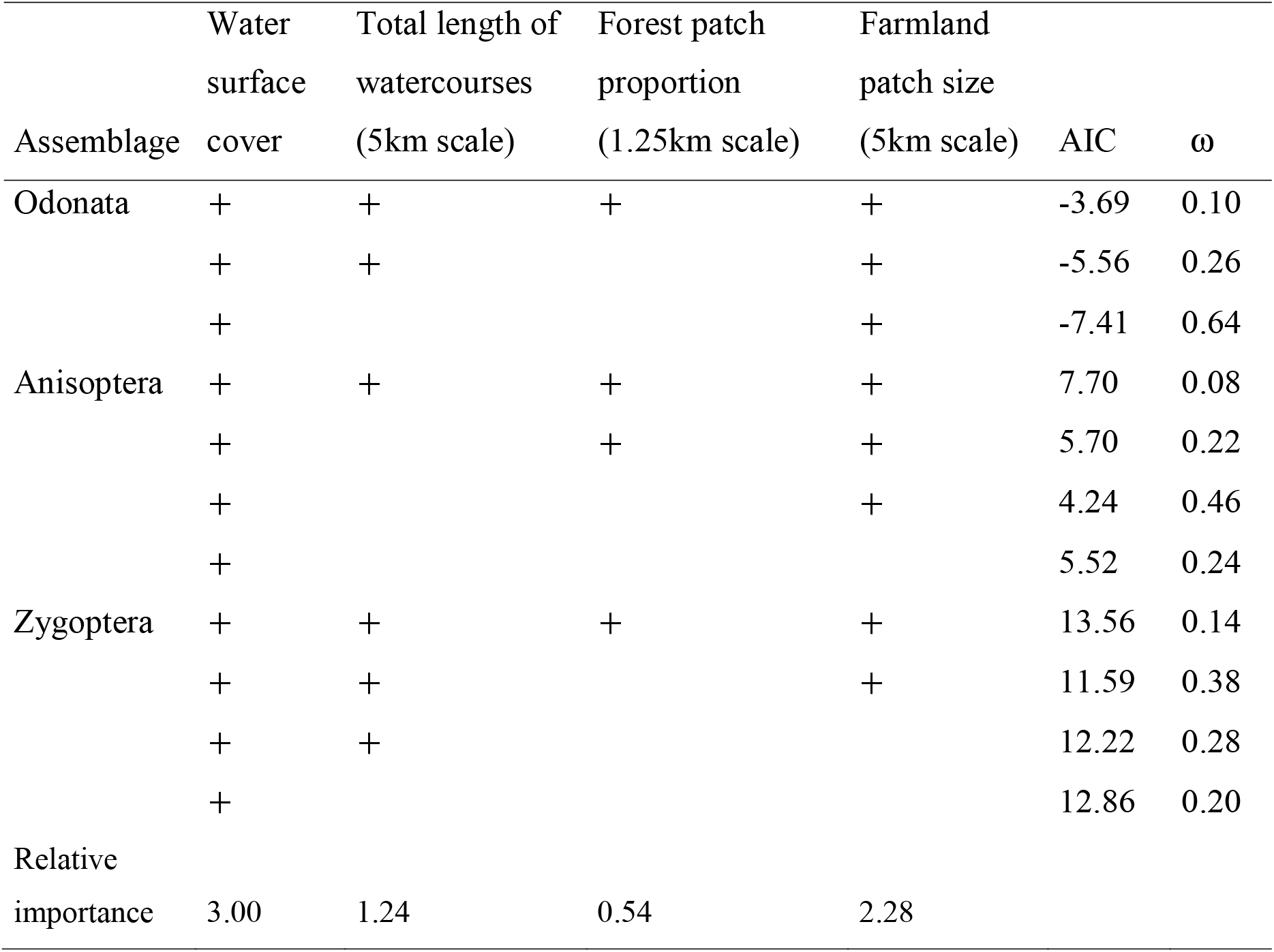
The analysed models (Gaussian errors) explaining species diversities of Odonata assemblages. AIC = Akaike’s information criteria. ω = Akaike weights. The “+” signs denote variables entered into the models.

## DISCUSSION

We tested the influence of water body attributes and the surrounding landscape on the Odonata assemblages along lowland watercourses. The hypothesis was that the studied watercourses were the most stable aquatic environments from the whole hydrographic basin, because they persisted even in extreme dry summer periods. The short term stability of the aquatic ecosystems is of crucial importance for the breeding success and generational continuity of populations in most of species. ^27^. Furthermore, the quality of the terrestrial environment is also important for the local populations because it provides habitats for mating, egg laying, feeding, resting and facilitates dispersal ^28^. Our findings show that both local and landscape variables are important for the presence and abundance of Odonata. However, the two groups showed different sensitivities to the local and landscape variables.

Only one out of the six analysed local variables, the cover of emergent vegetation had significant influence on the species richness of Odonata. Especially the Zygoptera showed significant sensitivity to this variable. Furthermore, Zygoptera diversity decreased with the increase of the cover of emergent vegetation. However, there was a relatively high cover of emergent vegetation on almost all sites. Such a high open water surface cover may hamper the movement of Zygoptera, which has weaker flying and dispersal ability than the Anisoptera species ^2^. Rouquette and Thompson ^19^ report the importance of emergent vegetation in the case of *Coenagrion mercurial*e; they underline that high percentage of water surface cover is not favoured, at the same time open water affects positively the density of *C. mercuriale*. In another study, where both Zygoptera and Anisoptera species were analysed from the perspective of water surface cover, anisopteran species were more affected than zygopteran ones ^18^. The explanation for the inconsistence between our results and the previously cited one is regarded to water surface cover variability. Most of the Zygoptera are perchers, and detect intruders, or females sitting on different surfaces (plants, sticks) by watching around ^2^. In our case the water surface cover was rather high which may prohibited the movement of Zygoptera.

The relationship between the landscape variables and Odonata was significant only for the Anisoptera species, and this result supported our expectation. As expected from published literature Anisoptera due to their higher dispersal ability are more sensitive to the landscape structure than Zygoptera. This assumption was also confirmed in other studies, where it was reported that the more mobile Anisoptera were more sensitive to landscape variables at large scales, while Zygoptera were sensitive to local (i.e. water body related) variables ^4,10^.

This difference in habitat sensitivity between the two Odonata suborders can explain the result that total length of watercourses on a 5 km scale has a positive significant effect on Anisoptera diversity. Anisoptera have larger size, bigger muscle mass, and better thermoregulation than Zygoptera ^29^, and thus better flying abilities. A longer watercourse network provides an extended habitat which means more food, more oviposition site and more conspecific females, and higher survival chance. In England Raebel et al. ^28^ found that the number of ponds in the surrounding area had no effect on species richness of dragonflies. However, their largest spatial scale was of 1600 meter long radius, contrary to the 5 km long radius scale used in our study. In another study ^28^ authors found that the distance to the nearest possible pond is a crucial factor in species occurrence: species richness decreased with increasing distance to the nearest suitable pond. In an experimental study where cattle tanks were used the results show that both the distance to the nearest tank and the connectivity between artificial ponds affected significantly the species richness ^15^. With increasing isolation the dispersal between tanks decreased, and thus species richness declined.

The farmland patch size showed a significant negative effect on Odonata species diversity at large scale (5 km), and on Anisoptera species diversity at the small (1.25 km) scale. The trend was the same for Anisoptera at the middle scale (2.5 km). The farmland patch size alludes to landscape fragmentation: increasing patch size results in landscape homogenization, with fewer buffer strips, bushes, forest patches, and presumably high fertilizer input. In the agriculturally intensified landscapes this means less space for maturation, feeding, and resting for the dragonflies.

The negative effects of the intensified land use on a large number of Odonata was presented by Ott ^30^ (1995). In another study on odonate species richness a similar effect was described where the species richness increased with larger areas of land under Higher Level Scheme ^28^. The Higher Level Scheme, an agri-environmental scheme include pond-specific options that could potentially beneficial for Odonata, by assuring buffering in-field ponds in improved grassland or farmland, maintenance of high quality ponds, and pond creation and restoration.

Habitat structure and landscape configuration effect on species diversity was demonstrated in a study (Georgian Bay region, Canada). They showed that the habitat structure and other landscape variables calculated at variable scales (at 1, 2, 4 and 8 km) was more important than boating pressure both for adults and for larvae ^31^. In a study of the threatened dragonfly species *Sympetrum depressiusculum* Dolný et al. ^32^ and Hykel et al. ^33^ suggest that the heterogeneous terrestrial habitat structure is essential for the development of juveniles, and movement of adults, which preferred habitat patches with abundant vegetation. When the importance of land cover types per se and landcover heterogeneity was studied, authors showed that from nine land cover types, farmland percentage had positive effect on 9 species, and negative effect on 31 species. They also found that in the case of 73 species abundance increased with the increasing of landcover heterogeneity ^21^.

Species diversity of Zygoptera showed a marginally significant decreasing trend with the increasing forest patch proportion in the surrounding habitat at the small scale (1.25 km). This relationship was underlined in a study where Odonata species richness decreased with increasing amounts of forest, especially at a 200 m scale ^28^. Although the role of forests for Odonata has a great literature ^5,14,34^ our result shows that for the lowland Zygoptera species the increased amount of woodland could be an obstructive factor.

We demonstrated that Odonata show different responses to local and landscape variables. While the Zygoptera were mostly affected by local variables, the Anisoptera were more sensitivity to landscape variables. Our study further highlights the need for simultaneous consideration of local (aquatic habitat related) and landscape variables to understand fully the habitat use of Odonata. Our findings suggest to use a management which support moderate vegetation cover and a heterogeneous, patchy vegetation. This kind of management support species-rich Odonata communities and may also be beneficial for several other taxa such as amphibians, butterflies and farmland birds.

## MATERIAL AND METHODS

### Sampling sites

Lowland watercourses and their neighbouring habitats of North East Hungary (Szatmári plain) and North West Romania (North-Partium) were surveyed in 2015 and 2016 (**Figure 2**). The region surveyed in Hungary had cca. 62 km^2^ and that of Romania had 127 km^2^. The two survey years were dry ^35^; in this context, we identified only 11 independent watercourses (4 watercourses in Hungary and 7 watercourses in Romania) with substantial amount of water in order to implement our sampling design. The chosen watercourses were characterised by the presence of *Carex* sp., *Glyceria maxima*, *Mentha aquatica*, *Nuphar lutea*, *Sparganium erectum*, *Stratiotes aloides*, *Phragmites australis*, *Typha latifolia*, *T. angustifolia*. Banksides had rich herbaceous vegetation, with scarce shrub and tree cover. Surveyed watercourses were at different distances from forest patches. All watercourses were at least on one bankside adjacent to agricultural fields, mostly farmlands. All surveyed watercourses were outside of urban areas.

**Figure 2.**
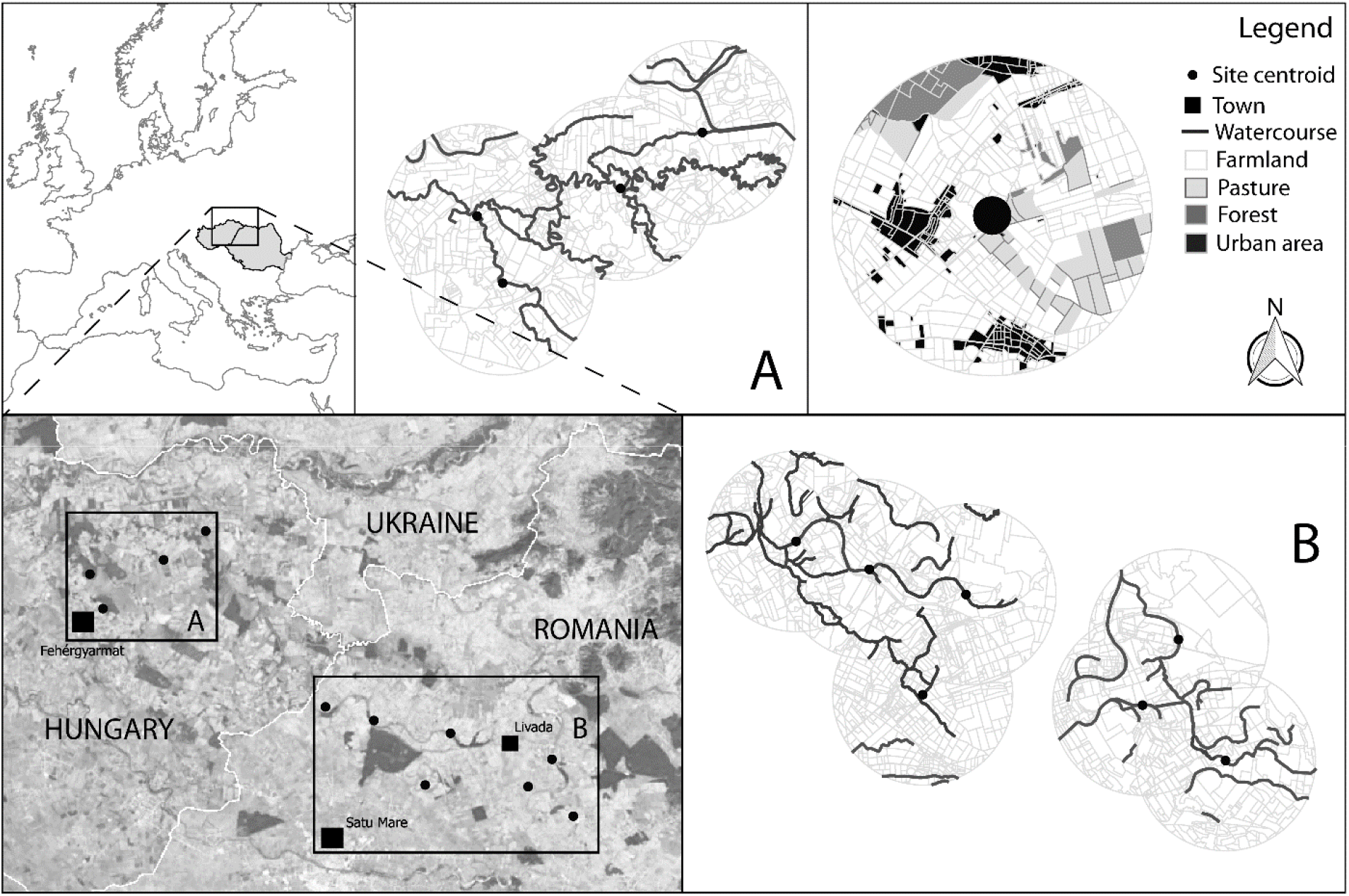
Sites and their landscape neighbourhood. The sampled watercourse segments are positioned at the site centroids. The predominant cover types were farmlands, urban areas, forest patches, and pastures. Cover type boundaries were manually digitised by the authors.

### Data collection

We sampled the Odonata assemblages using a 500-meter-long transect along each watercourse. The sampling events dated from May to September, once a month, in warm, sunny days when the minimum temperature exceeded 20°C, with wind speed under 15 km/h, and no considerable cloud cover were observed. The same person, walking at a steady pace counted every observed specimen in every sample event. Every specimen was identified to species level. Species were identified both either visually (e.g. this, that and that species) or were caught with an insect net when visual identification was not possible in other way (e.g. this, that and that species). Only those specimens were caught with an insect net, which were difficult to identify.

### Local and landscape variables

Local variables were recorded in six points across 500 m transect, for each studied watercourse. These were: water depth (meter, *m*), water width (*m*), macrovegetation cover (percentage cover), bankside cover with trees and bushes (percentage cover), bankside cover with herbaceous plants (percentage cover), and average height of the bankside vegetation (cm). These variables were measured or estimated by the same person at every single sample event. As local abiotic variables, we have measured once at every sample event the air temperature, wind speed, humidity, and the distance of visibility.

Landscape level variables were recorded in a circle with radii 1.25 (small scale), 2.5 (intermediate scale) and 5 km (large scale) around the midpoint of the sampled transects. These areas were digitised from the highest spatial resolution satellite images possible, acquired from Google Earth™ (http://earth.google.com/; © 2016 Google; © 2016 Geoeye; © 2016 DigitalGlobe). The maps were constructed from manually digitised cover type boundaries at a resolution ratio of 1:250 in Quantum GIS (version 2.14.11 “Essen”; Quantum GIS Development Team 2016). Cover types were delimited as farmland, pastures, orchards, urban areas, broad-leaved forests, bushy areas, embankments, dry riverbeds, rivers and lakes. The area (hectare) of each cover type was calculated using Quantum GIS.

From the digitised maps we calculated the following variables: landscape diversity with Shannon index, total length of watercourses, and proportion of forest patches, farmland patch size, and the distance to the nearest forest patch. We used the patch sizes of farmlands instead of their proportion in the landscape because the type of land use can be determined by the mean patch size of the crops in the landscape. All variables were calculated at all used scales, excepting the distance to the nearest forest patch, which was measured only in the smallest (1.25 km radius) circles. Total length of watercourses contained length of creeks and rivers. Landscape diversity, forest patch proportion, and farmland patch size were calculated using the package LecoS ^37^ in Quantum GIS.

### Statistical methods

The R programming language was used during the statistical calculations (R Development Core Team, version 3.5.0 2018). To assess Odonata assemblage characteristics we calculated Shannon diversity for each sampling site using function ‘diversity’ from package ‘vegan’ ^38^. We calculated Shannon diversity for Anisoptera, Zygoptera and all Odonata. Then we used Goodness of Fit (GOF) tests to verify the normality assumption for each analysed outcome variable. Collinearity between the explanatory variables was assessed with Pearson correlation; no collinearity was detected (*r*<0.5) therefore we used all variables in the modelling ^39^. For assessing correlations between local or landscape level environmental variables and assemblage diversities, we calculated Pearson’s correlation coefficients. We used those environmental variables that showed significant or marginally significant correlations to build linear models with Gaussian error distributions (using function ‘lm’). Backward stepwise selection procedure (using function ‘update’) were used selecting important variables. From all models, we calculated AICs and Akaike weights (ω) (using function ‘akaike.weights’ from package ‘qpcR’). Based on ω values we calculated the relative importance of the used environmental variable sets as described in ^39^.

## Acknowledgements

The Postdoctoral Fellowship Programme of the Hungarian Academy of Sciences funded the work of H. Beáta Nagy. Research were supported by OTKA K 116639, and KH 126477 projects. We thank the help during field work to Antal Orsolya-Mária and Lukács Alexandra. We are also indebted for the valuable comments and suggestions of Tibor Hartel.

## Compliance with ethical standards

The authors declare that they have no conflicts of interest.

## Author contributions

HBN and ZL initiated the project, collected data, interpreted results, and drafted the manuscript. FS and LS made landscape maps: digitized satellite photos. GD and BT initiated the project, contributed substantially to revisions. All authors gave final approval for publication.

