## Supplementary_material_Table_S1 for "Landscape scale terrestrial factors are also vital in shaping Odonata diversity of watercourses"

| **Table S1. Encountered specimen sums from years 2015, 2016 at each study site with site coordinates.** | | | | | | | | |  | |  | |  | |  | |  | |
| --- | --- | --- | --- | --- | --- | --- | --- | --- | --- | --- | --- | --- | --- | --- | --- | --- | --- | --- |
| **Species** | **Suborder** | **Site 1** | **Site 2** | **Site 3** | **Site 4** | **Site 5** | **Site 6** | **Site 7** | | **Site 8** | | **Site 9** | | **Site 10** | | **Site 11** | | **Total** |
| *Chalcolestes parvidens* | Zygoptera | 3 | 99 | 0 | 0 | 12 | 2 | 0 | | 5 | | 1 | | 3 | | 0 | | 125 |
| *Lestes barbarus* | Zygoptera | 0 | 219 | 2 | 12 | 0 | 0 | 0 | | 0 | | 0 | | 9 | | 1 | | 243 |
| *Lestes dryas* | Zygoptera | 0 | 0 | 0 | 0 | 0 | 0 | 0 | | 0 | | 0 | | 0 | | 0 | | 0 |
| *Lestes sponsa* | Zygoptera | 48 | 0 | 2 | 0 | 0 | 0 | 0 | | 0 | | 0 | | 0 | | 0 | | 50 |
| *Sympecma fusca* | Zygoptera | 33 | 3 | 0 | 4 | 1 | 0 | 0 | | 0 | | 2 | | 0 | | 0 | | 43 |
| *Calopteryx splendens* | Zygoptera | 2 | 52 | 107 | 4 | 59 | 5 | 21 | | 22 | | 1 | | 3 | | 71 | | 347 |
| *Platycnemis pennipes* | Zygoptera | 2 | 30 | 122 | 2 | 78 | 0 | 10 | | 3 | | 8 | | 1 | | 386 | | 642 |
| *Coenagrion puella* | Zygoptera | 263 | 180 | 171 | 8 | 283 | 127 | 46 | | 601 | | 20 | | 323 | | 846 | | 2868 |
| *Coenagrion pulchellum* | Zygoptera | 31 | 9 | 59 | 0 | 6 | 20 | 1 | | 53 | | 437 | | 3 | | 10 | | 629 |
| *Erythromma najas* | Zygoptera | 0 | 0 | 0 | 0 | 0 | 0 | 0 | | 0 | | 9 | | 0 | | 4 | | 13 |
| *Erythromma viridulum* | Zygoptera | 0 | 0 | 0 | 0 | 0 | 0 | 0 | | 0 | | 3 | | 0 | | 0 | | 3 |
| *Ischnura elegans* | Zygoptera | 10 | 2 | 165 | 14 | 36 | 3 | 0 | | 9 | | 5 | | 1 | | 41 | | 286 |
| *Ischnura pumilio* | Zygoptera | 6 | 0 | 35 | 24 | 140 | 4 | 2 | | 1 | | 8 | | 5 | | 1 | | 226 |
| *Pyrrhosoma nymphula* | Zygoptera | 0 | 4 | 0 | 0 | 0 | 0 | 1 | | 0 | | 0 | | 11 | | 0 | | 16 |
| *Aeshna affinis* | Anisoptera | 20 | 81 | 12 | 58 | 3 | 18 | 50 | | 34 | | 93 | | 26 | | 35 | | 430 |
| *Aeshna cyanea* | Anisoptera | 0 | 1 | 0 | 0 | 15 | 0 | 2 | | 0 | | 0 | | 2 | | 0 | | 20 |
| *Aeshna mixta* | Anisoptera | 5 | 3 | 3 | 0 | 0 | 2 | 0 | | 11 | | 33 | | 0 | | 5 | | 62 |
| *Anaciaeschna isoceles* | Anisoptera | 30 | 2 | 7 | 1 | 19 | 2 | 1 | | 15 | | 177 | | 2 | | 99 | | 355 |
| *Anax imperator* | Anisoptera | 7 | 0 | 4 | 2 | 0 | 0 | 1 | | 2 | | 2 | | 0 | | 1 | | 19 |
| *Brachytron pratense* | Anisoptera | 23 | 12 | 0 | 5 | 31 | 3 | 2 | | 0 | | 8 | | 3 | | 6 | | 93 |
| *Onychogomphus forcipatus* | Anisoptera | 0 | 0 | 0 | 0 | 0 | 0 | 0 | | 0 | | 1 | | 0 | | 0 | | 1 |
| *Somatochlora flavomaculata* | Anisoptera | 0 | 53 | 4 | 0 | 26 | 22 | 101 | | 0 | | 5 | | 0 | | 0 | | 211 |
| *Somatochlora meridionalis* | Anisoptera | 0 | 0 | 0 | 0 | 2 | 0 | 0 | | 0 | | 0 | | 0 | | 2 | | 4 |
| *Libellula depressa* | Anisoptera | 14 | 0 | 8 | 3 | 107 | 2 | 6 | | 7 | | 0 | | 20 | | 4 | | 171 |
| *Libellula fulva* | Anisoptera | 3 | 68 | 70 | 0 | 64 | 34 | 89 | | 6 | | 3 | | 0 | | 109 | | 446 |
| *Libellula quadrimaculata* | Anisoptera | 0 | 0 | 0 | 0 | 0 | 0 | 0 | | 0 | | 0 | | 0 | | 1 | | 1 |
| *Orthetrum albistylum* | Anisoptera | 0 | 0 | 2 | 2 | 0 | 0 | 0 | | 0 | | 0 | | 0 | | 0 | | 4 |
| *Orthetrum brunneum* | Anisoptera | 0 | 0 | 0 | 0 | 0 | 0 | 0 | | 3 | | 0 | | 0 | | 0 | | 3 |
| *Orthetrum coerulescens* | Anisoptera | 1 | 0 | 0 | 0 | 0 | 5 | 1 | | 0 | | 0 | | 0 | | 3 | | 10 |
| *Sympetrum flaveolum* | Anisoptera | 0 | 0 | 0 | 18 | 0 | 0 | 5 | | 0 | | 0 | | 0 | | 0 | | 23 |
| *Sympetrum meridionale* | Anisoptera | 90 | 22 | 0 | 70 | 0 | 0 | 0 | | 0 | | 0 | | 8 | | 16 | | 206 |
| *Sympetrum sanguineum* | Anisoptera | 105 | 1049 | 112 | 123 | 107 | 116 | 365 | | 142 | | 287 | | 191 | | 307 | | 2904 |
| *Sympetrum striolatum* | Anisoptera | 0 | 0 | 0 | 0 | 17 | 31 | 52 | | 66 | | 51 | | 41 | | 26 | | 284 |
| *Sympetrum vulgatum* | Anisoptera | 5 | 0 | 0 | 0 | 46 | 0 | 0 | | 2 | | 88 | | 0 | | 5 | | 146 |
|  | Country | RO | RO | RO | RO | RO | RO | RO | | HU | | HU | | HU | | HU | |  |
|  | E | 22,97974 | 23,01215 | 23,14433 | 22,93974 | 23,20644 | 22,88498 | 23,17116 | | 22,51728 | | 22,68234 | | 22,53596 | | 22,62234 | |  |
|  | N | 47,83253 | 47,8827 | 47,82729 | 47,89531 | 47,79951 | 47,90931 | 47,86018 | | 48,03023 | | 48,07118 | | 47,99738 | | 48,04377 | |  |
